# Poly-gamma-glutamic acid secretion protects *Bacillus subtilis* from zinc and copper intoxication

**DOI:** 10.1101/2021.08.18.456869

**Authors:** Reina Deol, Ashweetha Louis, Harper Lee Glazer, Warren Hosseinion, Pete Chandrangsu

## Abstract

Zinc and copper are essential micronutrients that serve as a cofactors for numerous enzymes. However, when present at elevated concentrations, zinc and copper are highly toxic to bacteria. To combat the effects of zinc and copper excess, bacteria have evolved a wide array of defense mechanisms. Here, we show that the Gram positive soil bacterium, *Bacillus subtilis*, produces the extracellular polymeric substance, poly-gamma-glutamate (γ-PGA) as a protective mechanism in response to zinc and copper excess. Furthermore, we provide evidence that zinc and copper dependent γ-PGA production is independent of the DegS-DegQ two component regulatory system and likely occurs at a post-transcriptional level. These data provide new insight into bacterial metal resistance mechanisms and contribute to our understanding of the regulation of bacterial γ-PGA biosynthesis.

**Importance:** Zinc and copper are potent antimicrobial compounds. As such, bacteria have evolved a diverse range of tools to prevent metal intoxication. Here, we show that the Gram-positive model organism, *Bacillus subtilis*, produces poly-gamma-glutamic acid (γ-PGA) as a protective mechanism against zinc and copper intoxication and that zinc and copper dependent γ-PGA production occurs by a yet undefined mechanism independent of known γ-PGA regulation pathways.

## Introduction

Transition metals ions, such as zinc and copper, are essential for life, yet toxic in excess. Metals have long been appreciated for their antimicrobial properties. Ancient Egyptians used copper salts as early as 2400 BCE as an astringent, food preservative, and disinfectant (1). Since zinc and copper disrupt antibiotic-resistant biofilms, display synergistic activity with other antimicrobials, and kill antibiotic-resistant bacteria, metal-based antimicrobials are currently used in industry, agriculture, and healthcare as metal-impregnated coatings and surfaces and in combination with traditional antibiotics (2). Furthermore, growing evidence suggest that zinc and copper intoxication play a key role in the host-microbe interaction as antibacterial agents (3).

At elevated concentrations, zinc and copper can outcompete other metals, including manganese and iron, for binding to metal containing proteins (4). Bacteria have evolved diverse mechanisms to prevent zinc and copper toxicity. Under zinc and copper excess, the intracellular level of these ions can be maintained by reduced uptake and/or increased efflux through the regulation of expression and activity of membrane associated metal transporters (5). Zinc and copper can also be detoxified by intracellular sequestration by small proteins or low molecular weight thiols, such as bacillithiol and glutathione (6, 7). The specific proteins and cellular processes poisoned by metals can also be repaired by chaperones or antioxidants, or bypassed through the expression of alternative metabolic pathways (8).

During the course of our investigations into the molecular mechanisms underlying bacterial zinc toxicity, we observed that *B. subtilis* colonies became highly mucoid when grown in the presence of zinc. Here, we demonstrate that the mucoid phenotype is due to the production of the secreted polymer, poly-gamma-glutamic acid (γ-PGA), show that γ-PGA protects *B. subtilis* from zinc and copper intoxication, and provide evidence that zinc and copper induce γ-PGA production independently from known regulatory mechanisms.

## Results

### Zinc and copper induce *B. subtilis* γ-PGA production

While studying the mechanisms of zinc intoxication, we observed that *B. subtilis* colonies displayed a mucoid phenotype (Figure 1A). *Bacillus subtilis* serves as a model organism for the study of biofilm formation and extracellular polymeric substance (EPS) production (9). There is a growing body of evidence suggesting a link between zinc and copper homeostasis and EPS production. In *B. subtilis*, zinc and copper decrease biofilm hydrophobicity, thereby increasing antibiotic effectiveness (10). Additionally, excess zinc decreases *Streptococcus pyogenes* hyaluronic acid capsule production by inhibiting phosphoglucomutase, a key enzyme in central carbon metabolism (11). Recent studies on *Escherichia coli* virulence suggest that the presence of capsule may exacerbate bile salt mediated zinc starvation (12).

**Figure 1:**
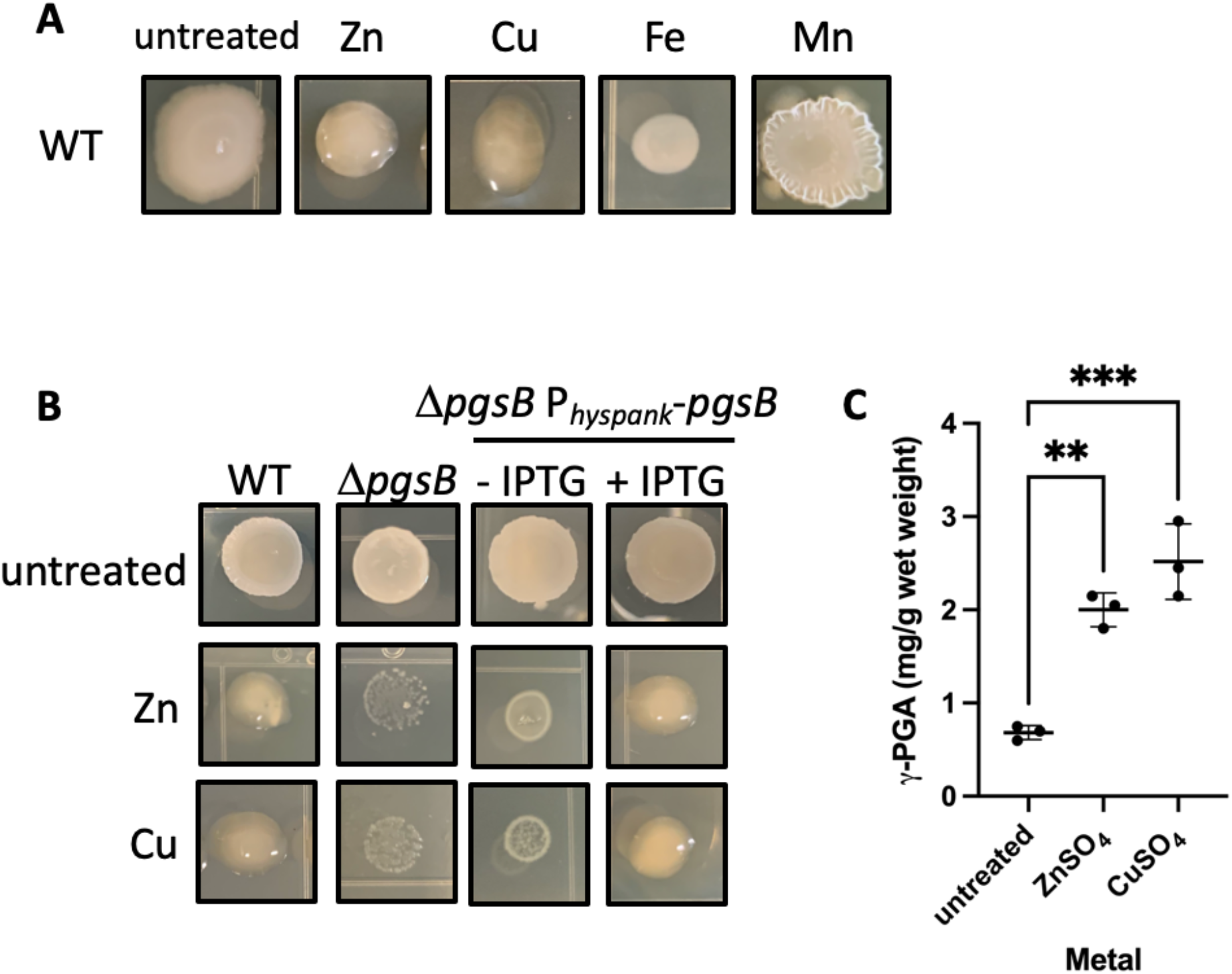
Zinc and copper induce PGA production. (A) Colony morphology of WT *B. subtilis* 3610 *comI* in the presence of 250 μM ZnSO_4_, 1 mM CuSO_4_, 2 mM FeSO_4_, 500 μM MnCl_2._ (B) Colony morphology of WT, Δ*pgsB*, and Δ*pgsB* P_hyspank_-*pgsB* grown on LB agar in the presence of 250 μM ZnSO_4_ or 100 μM CuSO_4_. IPTG (1 mM) was also added to Δ*pgsB* P_hyspank_-*pgsB* to induce *pgsB* expression. (C) Concentration of *γ-*PGA produced by cells grown in the presence of 250 μM ZnSO_4_ or 100 μM CuSO_4_. The *γ-*PGA concentration is expressed as mg of *γ-*PGA per gram of cell wet weight. Data shown are the mean and standard deviation from three independent experiments. *P* values were calculated by an ANOVA. **, *P* ≤ 0.01; ***, *P* ≤ 0.001

The EPS is composed of polysaccharides, proteins, nucleic acids, and poly-gamma-glutamic acid (γ-PGA) (13). γ-PGA is a biopolymer consisting of repeating units of L- or D-glutamic acid or a combination of both, and plays a role in virulence, survival during harsh conditions, and sequestration of toxic metal ions (14). In *B. subtilis*, the γ-PGA biosynthesis machinery is encoded by the *pgsBCAE* operon. In some bacteria, such as *B. subtilis*, γ-PGA is secreted and released from the cell surface, whereas in others like *B. anthracis* and *S. pneumoniae*, PGA remains attached to the cell as a capsule (15). When γ-PGA is made and secreted by *B. subtilis*, the colonies produce a mucoid slime layer. Therefore, we hypothesized that mucoid material is the result of γ-PGA production.

To determine if the mucoidy observed in the presence of zinc is due to increased γ-PGA production, we compared the colony phenotype of wild-type and a *pgsB* deletion mutant in the presence of zinc (Figure 1B). We observed that in the presence of a sublethal concentration of zinc (250 μM ZnSO_4_) wild-type colonies are mucoid and raised, whereas the *pgsB* mutant colonies are dull and flat. Furthermore, the mucoid phenotype is restored to the *pgsB* mutant when *pgsB* is expressed ectopically from an IPTG-inducible promoter (P_*hyspank*_).

To determine if the increase in γ-PGA production is specific to zinc or a general response to metals, we tested for colony mucoidy in the presence of a number of physiologically relevant transition metals (iron, manganese, or copper). Previously reported sublethal metal concentrations (2mM FeSO_4_, 500 μM MnCl_2_, or 1 mM CuSO_4_) were individually added to LB agar (16, 17). As previously reported, exposure to manganese leads to a wrinkled colony phenotype (18). Of the metals tested, only zinc and copper are able to induce colony mucoidy (Figure 1B) As observed with zinc, the colony mucoidy observed in the presence of copper is dependent on the presence of *pgsB*. Additionally, we utilized a spectrophotometric based assay to measure γ-PGA production in liquid media. Exposure to 250 μM ZnSO_4_ or 100 μM CuSO_4_ resulted in 3- or 5- fold increase in γ-PGA secretion (Figure 1C). From these data, we conclude that zinc and copper increase *B. subtilis γ-*PGA production.

### γ-PGA protects *B. subtilis* from zinc and copper intoxication

γ-PGA can directly bind to metals, such as zinc, copper, and silver, consistent with a proposed protective role for γ-PGA (19, 20). To determine if γ-PGA production serves as a means of protecting *B. subtilis* from high levels of zinc and copper, we compared the zinc and copper sensitivity of wild-type and a Δ*pgsB* mutant by a disk diffusion assay (Figure 2A). The Δ*pgsB* mutant is more sensitive to zinc and copper than wild-type. Furthermore, the zinc and copper resistance of the Δ*pgsB* mutant is restored when *pgsB* is expressed ectopically from an IPTG-inducible promoter.

**Figure 2:**
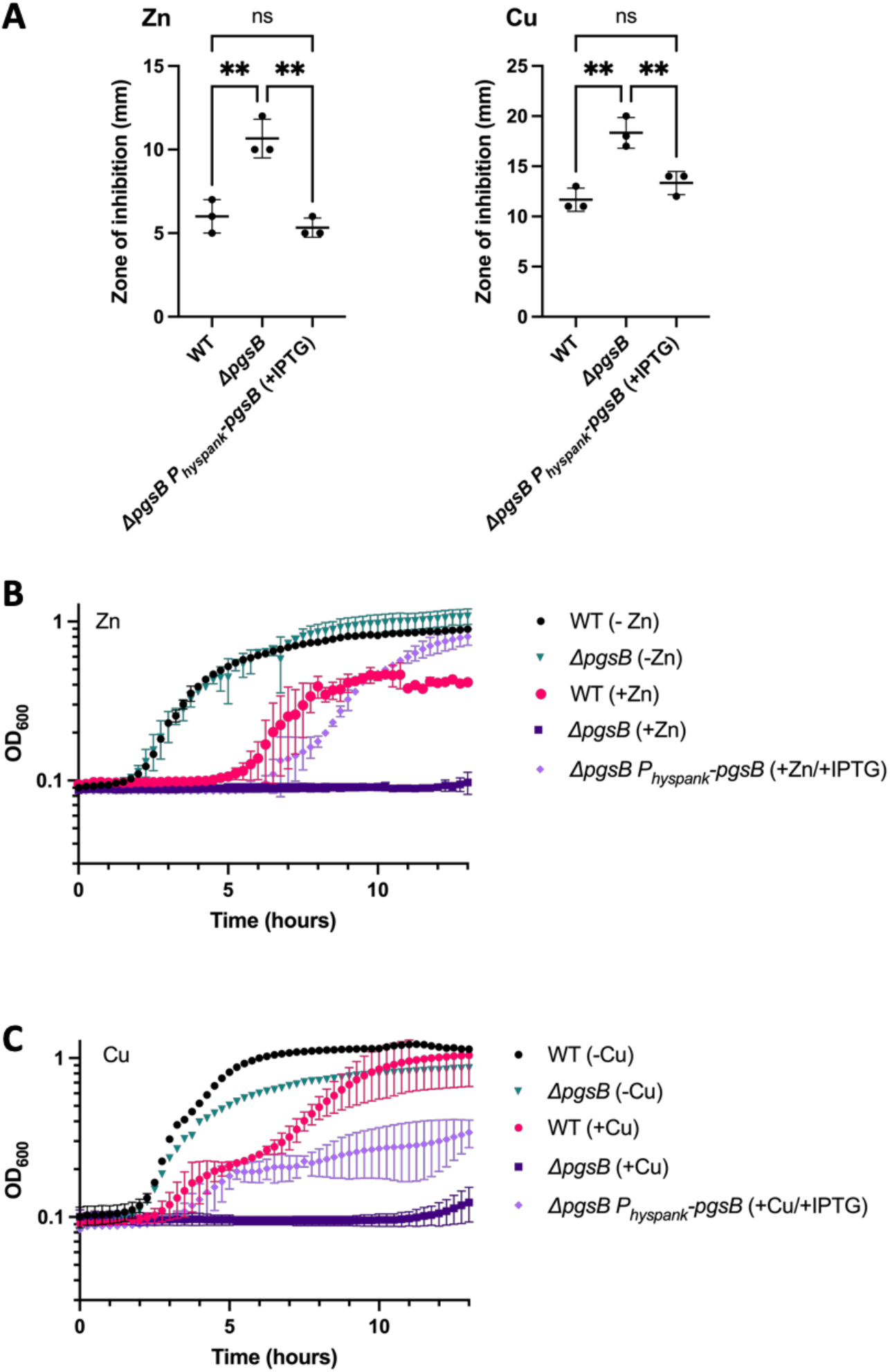
PGA protects *B. subtilis* from zinc and copper intoxication. (A) Disk diffusion susceptibility test performed on WT, Δ*pgsB*, and Δ*pgsB* P_hyspank_-*pgsB* grown on LB agar. Five μl of 500 μM ZnSO_4_ or 5 μl of 100 μM CuSO_4_ was added to a filter disk. The zone of inhibition was determined by measuring the diameter of the zone of clearing. Data shown are the mean and standard deviation from three different experiments. *P* values were calculated by an ANOVA. n.s,. nonsignificant; **, *P* ≤ 0.01. (B and C) Growth curves of WT, Δ*pgsB*, and Δ*pgsB* P_hyspank_- *pgsB* in the presence and absence of either (B) 250 μM ZnSO_4_ or (C) 100 μM CuSO_4_. IPTG (1 mM) was also added to Δ*pgsB* P_hyspank_-*pgsB* to induce *pgsB* expression. Data shown are the mean and standard deviation from three independent experiments.

We also assessed the contribution of γ-PGA to zinc and copper resistance by monitoring cell growth in LB broth in the presence or absence of zinc or copper (Figures 2B and C). Wild-type growth is impared in the presence of 250 μM ZnSO_4_ or 100 μM CuSO_4_ as indicated by an extended lag phase, decreased growth rate or decreased final OD_620_. In the absence of metal, growth of the *pgsB* mutant is similar to wild-type. Upon the addition of zinc or copper, the *pgsB* mutant is unable to grow. When *pgsB* is expressed from an IPTG-inducible promoter, growth of the *pgsB* mutant in the presence of zinc or copper increases. Restoration of growth is to near wild type levels in the case of zinc. In the presence of copper, wild-type like growth is restored during exponential phase. However, the *pgsB* complemented strain is unable to achieve the same final OD_600_ as wild-type. Since metals are involved in numerous cellular processes, the partial rescue in the presence of copper may be indicative of differences is the physiological processes inhibited by zinc and copper. From the disk diffusion and growth data, we conclude that γ-PGA protects *B. subtilis* from zinc and copper intoxication on both solid and liquid media.

### Zinc and copper dependent γ-PGA production is not controlled at the level of transcription

γ-PGA production can be regulated by altering the expression of the *pgs* operon, which encodes the PGA biosynthetic machinery, or of *pgdS*, which encodes a gamma-DL-glutamyl hydrolase involved in γ-PGA degradation (21, 22). Activation of *pgsB* operon expression involves the DegS-DegU two-component system, the ComP-ComA quorum sensing system, and the SwrA protein. (23). DegS-DegU control the expression of genes involved in many cellular behaviors, including biofilm formation, motility, and competence (24). In response to a variety of stimuli, the cytoplasmic DegS sensor histidine kinase phosphorylates the DegU response regulator (DegU-P), while DegQ enhances DegS-dependent phosphorylation of DegU (23, 25). Once phosphorylated, DegU-P activates *pgs* operon expression by binding upstream of the *pgsB* promoter (26, 27). ComP-ComA and SwrA indirectly activate *pgsB* operon expression by modulating DegU phosphorylation, in the case of SwrA or by regulating DegQ expression, in the case of ComP-ComA (27–29). Since DegS-DegU is downstream of ComA-ComP and SwrA, we decided to focus on DegS-DegU for further study.

To test if DegU is involved in the zinc or copper dependent production of PGA, we monitored the colony morphology of a *degU* mutant on LB agar in the presence of zinc or copper (Figure 3). Similar to wild-type, the *degU* mutant forms a highly mucoid colony in the presence zinc or copper. We infer that DegU is not required for zinc or copper dependent PGA production.

**Figure 3:**
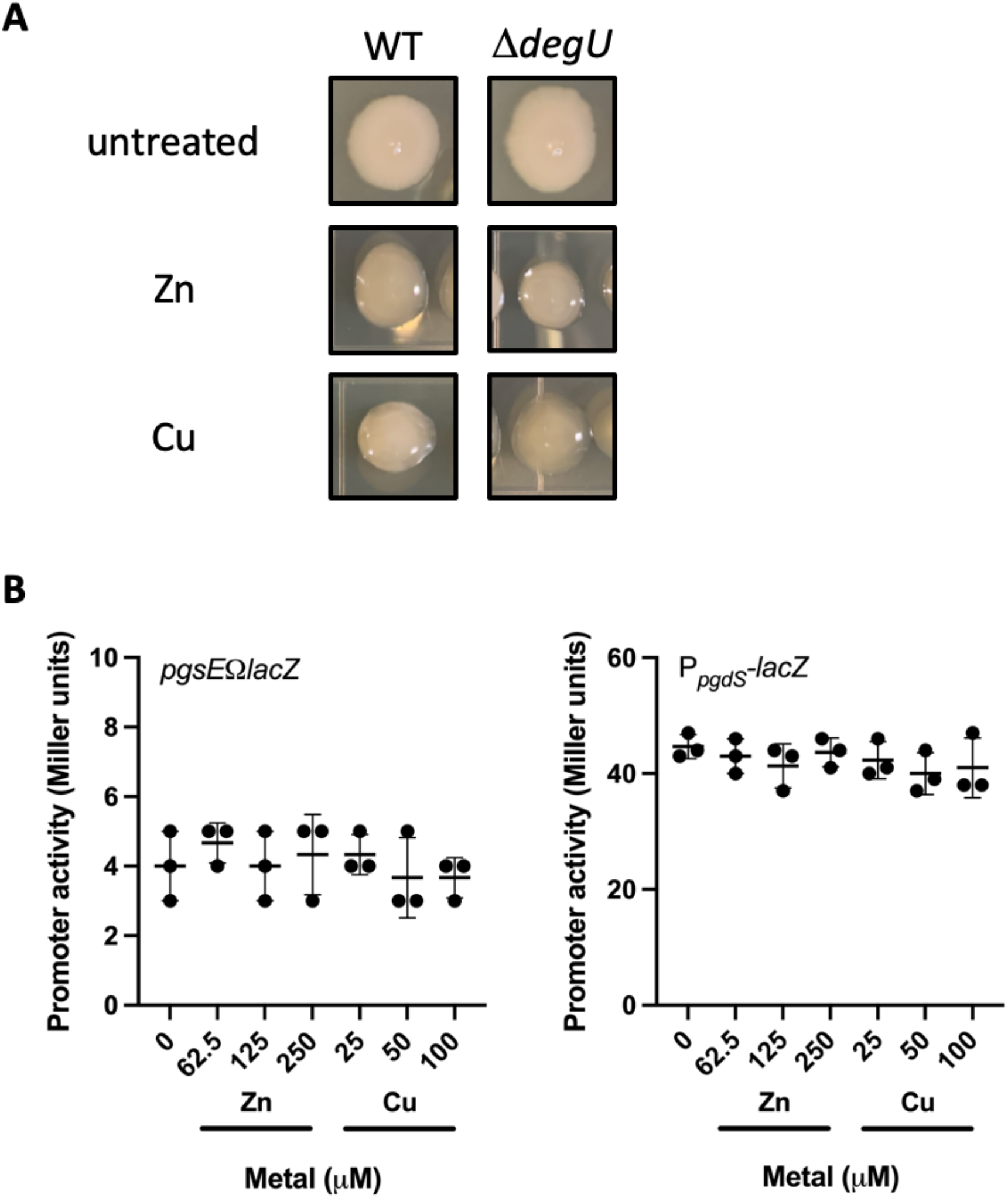
Zinc and copper dependent PGA production is not controlled at the level of transcription. (A) Colony morphology of WT and Δ*degU* grown in LB agar in the presence of 250 μM ZnSO_4_ or 1 mM CuSO_4_. (B) *pgs* operon (*pgsEΩlacZ*) and P_*pgdS*_ promoter activity in cells grown in LB media amended with varying concentrations of ZnSO_4_ or CuSO_4_. Data shown are the mean and standard deviation from three independent experiments. No significant difference was detected between samples by ANOVA.

We next explored the possibility *pgs* operon or *pgdS* expression may be regulated in response to zinc or copper excess. To determine if expression of the *pgs* operon or *pgdS* is differentially regulated in response to zinc or copper, we used a β-galactosidase assay to measure *pgs* operon and *pdgS* promoter activity in the presence of increasing concentrations of zinc or copper (Figure 3B). Even when grown in the presence of zinc (250 μM) and copper (100 μM) concentrations known to stimulate γ-PGA production, pgs operon and *pgdS* promoter activity did not substantially change. We conclude that zinc and copper dependent γ-PGA production does not occur through changes at the level of *pgs* or pgdS gene expression.

## Discussion

Our studies are consistent with previous work showing that *B. subtilis* produces γ-PGA in response to excess zinc. We extend these previous studies to demonstrate that γ-PGA production is also induced by copper. We demonstrate that γ-PGA provides *B. subtilis* protection from zinc and copper intoxication to provide a more complete picture of bacterial metal resistance.

Since γ-PGA is a non-toxic, biodegradable compound with utility in medical and commercial applications, microbial biosynthesis of γ-PGA is an extensively studied process (30, 31). Given its anionic charge and affinity for metals, γ-PGA is particular interest as a tool for the bioremediation of toxic metal ions from contaminated water (32). Thus, there is great interest in optimizing growth conditions and understanding the genetic circuitry underlying γ-PGA production. Based on our results, the mechanism of zinc and copper induction of γ-PGA production is independent of the DegS/U two component system known to activate *pgs* operon expression. Furthermore, our data suggest that regulation may be post-transcriptional, since neither γ-PGA biosynthesis (*pgsBCAE*) nor degradation (*pgdS*) gene expression is affected by the presence of excess zinc or copper.

Future investigations will explore the post-transcriptional regulation of γ-PGA biosynthesis. An intriguing candidate is PgsE, a small, 55 amino acid protein encoded last gene in the *pgs* operon. The *B. anthracis* PgsE homolog (CapE) is required for γ-PGA synthesis and is believed to play a structural role (33). Interestingly, when PgsE is overexpressed from an inducible promoter on high copy plasmid in the presence of zinc, γ-PGA production by *B. subtilis (chungkookjang)* increases 3-fold compared to the absence of PgsE overexpression (34). However, its role under physiologically relevant conditions is unclear. PgsE (*B. anthracis* CapE) is thought to associate with the PgsBCA (*B. anthracis* CapBCA) biosynthetic machinery in *B. anthracis* and when heterologously expressed in *E. coli* (33, 35). It is tempting to speculate that zinc and copper may be modulating PGA levels by affecting the interaction between PgsE and the PgsBCA protein complex. In total, our continuing work will provide further insight into the molecular mechanism underlying zinc and copper dependent γ-PGA production.

## Materials and Methods

### Strain and growth conditions

Strains were grown in Luria-Bertani broth (10 g tryptone, 5 g yeast extract, 5 g/l NaCl) or plates containing 1.5% agar at 37°C. When necessary, antibiotics were added to the growth media at the following concentrations: 1 μg/ml erythromycin plus 25 μg/ml lincomycin (mls), 100 μg/ml spectinomycin, 5 μg/ml kanamycin, 5 μg/ml chloramphenicol or 100 μg/ml ampicillin. One mM of isopropyl β-D-thiogalactopyranoside (IPTG) was added to the growth medium when appropriate.

### Strain construction

All strains and primers used in this study are listed in Tables 1 and 2, respectively. To facilitate genetic manipulation, all experiments were performed using the highly competent derivative of *Bacillus subtilis* 3610 carrying a *comI* deletion (3610 *comI*) (36). Chromosomal DNA was used for transformation as previously described (37). To induce natural competence, cells were grown in minimal competence media (1.7 g K_2_HPO_4_, 0.52 g KH_2_PO_4_, 2 g dextrose, 0.088 g sodium citrate dehydrate, 0.2. g l-glutamic acid monopotassium salt, 1 ml of 1,000× ferric ammonium citrate, and 1 g/l casein hydrolysate and 1% 300 mM MgSO_4_.

### Colony mucoidy assay

To evaluate the colony mucoidy phenotype, strains were grown to mid-exponential phase (OD_600_∼0.4) in LB broth Five μl of the culture were then spotted on to freshy poured LB plates containing 1.5% agar that had been dried in the laminar flow hood for 10 minutes. The spots were allowed to dry an additional 10 minutes in the laminar flow hood. The plates were then incubated at 37°C for 16-18 hours. Images of the colonies were recorded using an iPhone 12 held in a tripod 36 inches from the agar surface.

### PGA quantification

*B. subtilis* PGA production was measured using a cetryltrimethylammoonium bromide (CTAB) assay as previously described (38, 39). Cells were grown in LB broth containing various concentratrations of ZnSO_4_ or CuSO_4_ in LB media overnight at 37°C for 16-18 hours. Cells were then pelleted from 15 ml of culture by centrifugation at 10000 x g for 30 minutes at 4°C and the supernatant was retained. To precipitate the PGA, thirty ml of ethanol (96%) was added to the supernatant and incubated at 4°C overnight. The precipitated PGA was harvested by centrifugation at 10000 ⨯ g for 10 minutes at 4°C. The resulting pellet was resuspended in 1 ml of distilled water. In a 96 well flat bottom microtiter plate, one hundred μl of 0.07 M CTAB in 2% (w/v) NaOH was added to 100 μl of the resuspended PGA and mixed gently by pipetting to avoid bubble formation. The OD_400_ of the resulting solution was measured in a Tecan Sunrise microplate reader. A standard curve using purified PGA (Sigma, G1049) was used to calculate the PGA concentration in the broth.

### *pgsB* complementation

The plasmid pDR111 was used to generate the IPTG-inducible *amyE::P*_*hyspank*_*-pgsB* construct. Briefly, a PCR product containing the *pgsB* coding region was amplified from *B. subtilis* 3610 *comI* chromosomal DNA using primers 1003/1004, digested with SalI and SphI, and cloned into the SalI and SphI sites of the pDR111 plasmid. The plasmid was then transformed in to *B. subtilis* 3610 *comI*. Successful transformation was indicated by spectinomycin resistance and verified by PCR.

### In frame markerless gene deletions

Gene deletion mutants were constructed from the BKE strains as previously described (40). BKE strains carrying the gene deletion of interest marked by a kanamycin or erythromycin resistance cassette were acquired from the Bacillus Genetic Stock Center (www.bgsc.org). Chromosomal DNA was isolated and transformed in to our wild-type 3610 *comI* strain. To remove the antibiotic resistance cassette, the temperature sensitive pDR244 plasmid encoding the *cre* recombinase was introduced into the transformants by selecting for spectinomycin resistance at 30°C. The strains was subsequently cured of the plasmid by growth at 42°C and screened for sensitivity to spectinomycin (loss of the plasmid) and kanamycin or erthyromycin (loss of the antibiotic resistance cassette). Successful gene deletion was verified by PCR using primers 1001/1002.

### β-galactosidase assay

A β-galactosidase assay was used to measure *pgs* operon and P_*pdgS*_ promoter activity, as previously described (41). Cells were grown in LB broth in the presence or absence of ZnSO_4_ (62.5, 125, or 250 μM) or CuSO_4_ (25, 50, 100 μM) at 37°C. One ml of cells were harvested by centrifugation at early stationary phase (OD_600_∼1) and resuspended in an equal volume of Z-buffer (40 mM NaH_2_PO_4_, 60 mM Na_2_HPO_4_, 1 mM MgSO_4_, 10 mM KCl, and 38 mM 2-mercaptoethanol. Cells were lysed by addition of lysozyme (0.2 mg/ml) and incubation at 30°C for 15 minutes. Each sample was then diluted in Z-buffer to a final volume of 500 μl. The reaction was started with the addition of 100 μl of 4 mg/ml of 2-nitrophenyl β-D-galactopyranoside (ONPG). Once the sample began to turn yellow, the reaction was stopped with 250 μl of 1 M Na_2_CO_3_. The OD_420_ was measured, and the specific activity of β-galactosidase was calculated using the equation [OD_420_/(reaction time x OD_600_)] ⨯ dilution factor ⨯ 1000.

### Growth curves

To measure the effect of zinc and copper on cell growth, strains were grown to mid-exponential phase (OD_600_∼0.4) and subsequently diluted 1:100 into fresh LB media in the presence or absence of ZnSO_4_ (250 μM) and CuSO_4_ (100 μM) in a 96 well flat bottom microtiter plate. The plate was then incubated in a Tecan Sunrise microplate reader at 37°C with continuous shaking (high setting) and the OD_620_ was measured every 15 minutes for 16 hours.

### Disk diffusion assay

Disk diffusion assays were modified from (42). Briefly, a 1:100 dilution of an overnight culture was grown in fresh LB medium at 37°C with 250 rpm shaking until mid-exponential phase (OD_600_∼0.4). A 100 μl aliquot of cells was then spread on the surface of a LB agar plate (15 ml of 1.5% agar) that had been dried in a laminar flow hood for 10 minutes. The inoculum was allowed to dry for an additional 10 minutes in the laminar flow hood. A sterile 6.5 mm Whatman filter disk was then placed on the surface of the agar and ZnSO_4_ (5 μl of a 500 mm solution) or CuSO_4_ (5 μl of a 100 mm solution) was added to filter disk and allowed to absorb into the disk for 5 minutes. The plates were incubated at 37°C for 16-18 hrs, after which the diameter of the zone of inhibition was measured.

## Acknowledgements

We would like to thank Daniel Kearns (Indiana University) for insightful discussion and generously providing strains.

## Strains used in this study

| Strain | Description | Source |
| --- | --- | --- |
| PC1 | 3610 *comI* (wild-type) | (36)(36)(36) |
| BKE35900 | *trpC2* Δ*pgsB::erm* | (40) |
| PC2 | PC1 Δ*pgsB* | This work |
| PC3 | PC1 Δ*pgsB amyE::P<sub>hypspank</sub>-pgsB spec* | This work |
| BKK35490 | *trpC2 ΔdegU::kan* | (40) |
| PC7 | PC1 Δ*degU* | This work |
| DS9309 | 3610 *amyE::P<sub>pgdS</sub>-lacZ cat* | (41) |
| DS9079 | 3610 *pgsEΩlacZ cat* | (41) |
| PC10 | PC1 *amyE::P<sub>pgdS</sub>-lacZ cat* | This work |
| PC11 | PC2 *pgsEΩlacZ cat* | This work |

## Plasmids used in this study

| Plasmid | Description | Source |
| --- | --- | --- |
| pDR244 | *cre ori<sub>Ts</sub> amp spec* | (40) |
| pDR111 | *bla amyE' P<sub>hyperspank</sub> spec lacI 'amyE* | Gift from D. Kearns Lab |
| pPC1 | pDR111 *bla amyE' P<sub>hyperspank</sub> -pgsB spec lacI 'amyE* | This work |

## Primer table

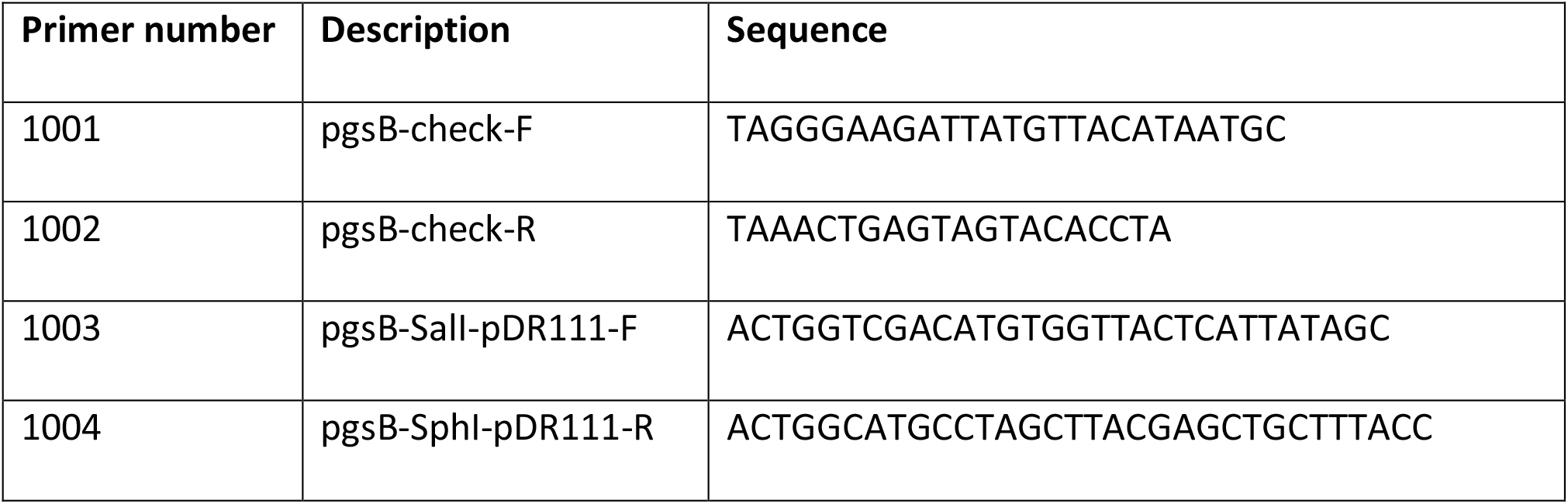

